# Universal inverse square relationship between heart rate variability and heart rate

**DOI:** 10.1101/2021.09.10.459420

**Authors:** Anna V. Maltsev, Oliver Monfredi, Victor A. Maltsev

**Author notes:** **Corresponding author:** Anna V. Maltsev, PhD.

## Abstract

In our previous study, we analyzed heart rate variability and heart rate from a large variety of cardiac preparations (including humans, living animals, Langendorff-perfused isolated hearts, and single sinoatrial nodal cells) in diverse species, combining our data with those of previously published articles. The analysis revealed that regardless of conditions, heart rate variability (for the purposes of the study assessed as standard deviation of beat-to-beat intervals) vs. heart rate follows a universal exponential decay-like relationship. Numerical simulations of diastolic interval variability by adding a randomly fluctuating term (I_per_) to net current revealed a similar relationship. In the present study, using a Taylor series, we found that this relationship is, in fact, inverse square, and we derive an explicit formula for the standard deviation (sd) of the cycle length (CL) as a function of heart rate (HR) with biophysically meaningful parameters: sd(CL)=sd(I_per_)*(60,000/mean(HR) -APD)^2/(ΔV*C), where CL is in ms, HR in beats per minute, I_per_ in pA, APD in ms is an average AP duration of pacemaker cells, C in pF is cell membrane capacitance, and ΔV is the magnitude of diastolic depolarization in mV. This relationship gives direct insight into heart rate variability mechanisms at the basic level of individual pacemaker cells, i.e. their intrinsic CL variability linked to stochastic operation of ion channels (both Ca release and cell membrane channels) generating I_per_. Our explicit formula may be also used for a more precise biomedical interpretation of heart rate variability after respective corrections for heart rate.

---

Indices of heart rate variability (*HRV*) are frequently used to predict cardiovascular and overall risk in both healthy populations and in those with disease. This is because *HRV* in part reflects autonomic nervous signaling to the heart. *HRV* changes substantially with aging and under different physiological, pathological and experimental conditions. However, to date, extensive research has failed to reliably and reproducibly extract usable information about health status from *HRV* data, especially in older patients and those with cardiovascular disease. This is reflected in the fact that few, if any, *HRV* indices find themselves being utilized in routine clinical practice. In general, the problem is that *HRV* has a highly complex nature, being generated from integrated signaling at different scales, from the subcellular level, to cells, to organs, to organ systems and organisms (including critical and incompletely understood heart-brain-vascular interactions). Few cardiovascular experts would dispute that *HRV* and its analysis hold important information about both intrinsic and extrinsic mechanisms influencing the heartbeat, and when carefully handled these could accurately predict and detect both the healthy and diseased state. However, deciphering and harnessing the *HRV* for practical broad clinical use remains a major biomedical challenge. One specific problem that we study here is that *HRV* and heart rate (*HR*) are intrinsically related, and as such the *HR* at which *HRV* was measured has to somehow be taken into account before any firm conclusions are drawn from the *HRV*. Our work aims at understanding how.

In our previous study^1^, we analyzed *HRV* and *HR* from a large variety of cardiac preparations (including humans, living animals, Langendorff-perfused isolated hearts, and single sinoatrial nodal cells) in diverse species, combining our data with those of previously published articles. The analysis revealed that regardless of conditions, *HRV* (for the purposes of the study assessed as standard deviation, or *sd*) of beat-to-beat intervals (or cycle length, *CL*) and *HR* follows a universal exponential decay-like relationship. At a basic level, *HRV* emerges as an intrinsic property of individual pacemaker cells *via* variability of the diastolic interval (*DI*)(Fig.1A). All subsequent autonomic signaling is superimposed onto this. Numerical simulations of *DI* variability by adding a randomly fluctuating term, perturbation current (*I*_*per*_), to net current revealed a similar relationship^1^.

In the present study, using a Taylor series, we found that this relationship is, in fact, inverse square, and we derive an explicit formula for *sd*(*CL*) *vs. HR* with biophysically meaningful parameters. This relationship is equivalent to the square root relationship of 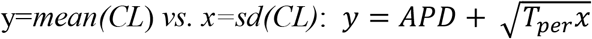, where *T*_*per*_ is a new parameter (dubbed “perturbation time”) given by 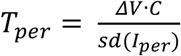, representing the time required for a membrane current with magnitude of *sd(I*_*per*_) to fully account for the diastolic depolarization amplitude Δ*V*=*E*_*th*_*-MDP*, and *C* is cell membrane capacitance (see derivation below).This relationship has units ms vs. ms and is thus easier to derive and interpret, whereas *sd(CL)* vs. *HR* is explicitly non-linear because of different units, ms vs. bpm.

We then fit our new theoretical function to the entire collection of experimental data from Monfredi et al.^1^ obtaining a correlation coefficient of 0.84, *APD*=131 ms, and *T*_*per*_=5885ms (Fig. 1B). The fitted *APD* is close to that in humans (∼140 ms), supporting our theory. In turn, *T*_*per*_ is directly linked to *I*_*per*_, i.e the intrinsic *CL* perturbation source (such as stochastic opening/closing of ion channels during the *DI*, and the “Calcium clock”^3^). Thus, *T*_*per*_ may have a new clinical and fundamental importance. Altogether, *mean(CL)* equals the sum of *APD* and a geometric average of *sd(CL)* and *T*_*per*_. Using our fit, *sd(I*_*per*_) is estimated to be ∼0.1 pA which is reasonable for pacemaker cell biophysics, and constitutes about 5% of the net membrane current during *DI* in human SA node cells (∼2 pA^4^), which also matches *CL* variability at a few percent scale under normal conditions. Our results are also supported by recent data from rabbit sinoatrial nodal cells^5^ closely fitted (R^2^=0.98) to a power function with power of 0.44, which is remarkably close to 0.5, i.e. the square root relationship derived here.

**Figure 1.**
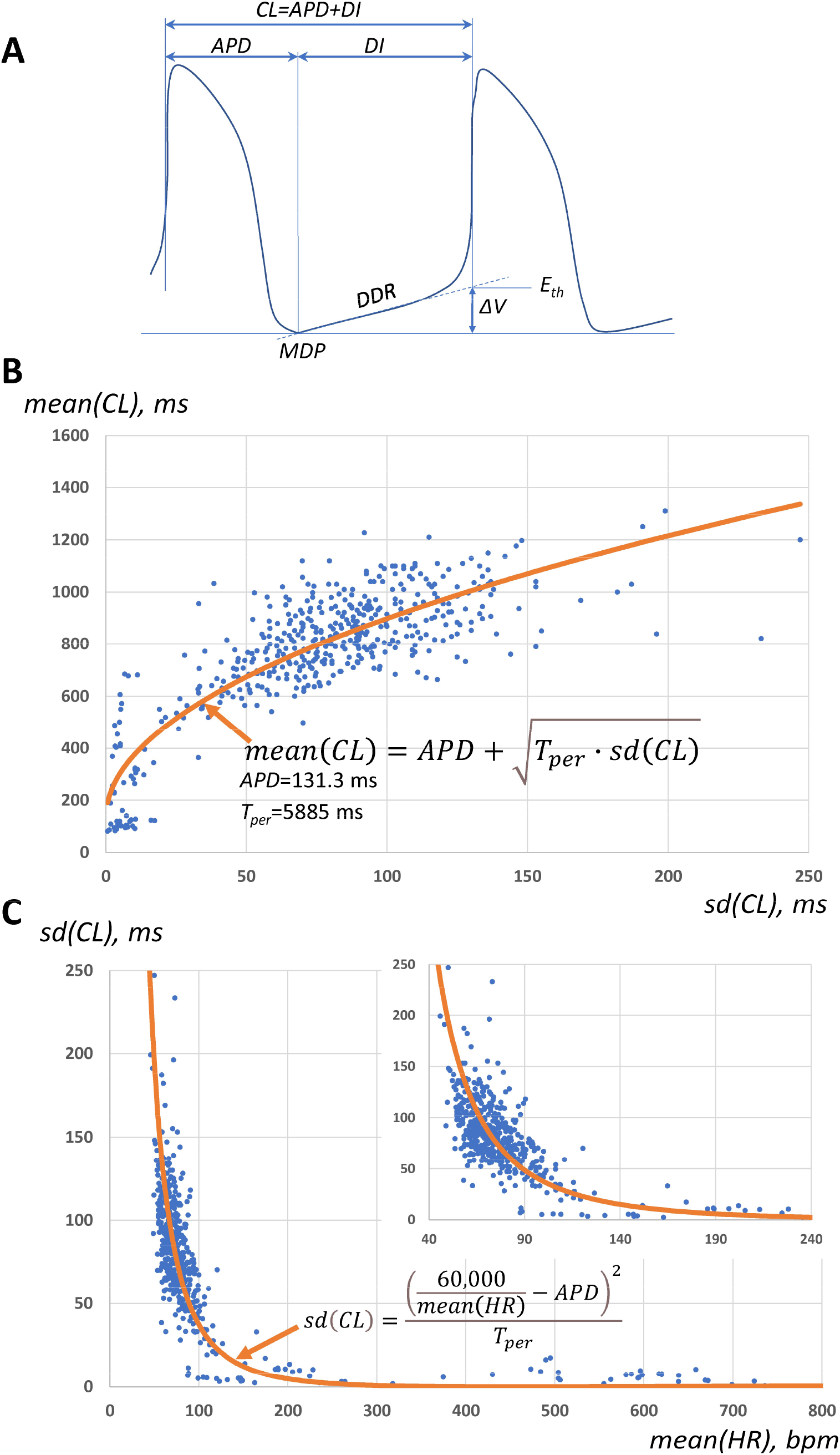
**(A)** Definitions used in the present study (introduced by Rocchetti et al.^2^): *CL*, cycle length; *APD*, AP duration, *DI*, diastolic interval; *E*_*th*_, AP threshold; *DDR*, diastolic depolarization rate; *MDP*, maximum diastolic potential. **(B and C)** Data from^1^ fitted with respective square root and reverse square relationships; inset shows the result within 40-240 bpm range (see text for details).

In conclusion, *HRV* is, in part, a non-invasive measure of autonomic tone to the heart, and can reflect patient morbidity and mortality. However, measures of *HRV* in different circumstances (e.g. males vs females, normal patients vs those with disease, athletes vs non-athletes etc) must to take account of the important intrinsic relationship between *HRV* and *HR* if the heart rate was different between the comparative groups. In our prior work^1^, the relationship was characterized in vague phenomenological terms as exponential decay-like relationship, which lacks both theoretical meaning and mechanistic interpretation. Here, we obtain this intrinsic relationship explicitly, with biophysically meaningful parameters, which will be of crucial importance for our further understanding of *HRV* mechanisms. Our explicit formula may be also of use for drawing biomedical conclusions from the *HRV* as it will allow for quantitatively accurate corrections for *HR*.

## Methods

Below is the full derivation of our formulas. Introducing average current *I* during *DI*, we obtain

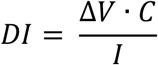

Suppose *I = I*_*0*_ *+ I*_*per*_, where < *I* >*= I*_*0*_ and *I*_*per*_ is a compactly supported random variable, <*I*_*per*_>*=0*, and *I*_*per*_^*2*^ is small. Using a Taylor expansion, we find

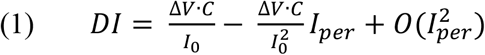

Taking mean and standard deviation of equation (1), and then noting that *CL=APD + DI*, we obtain

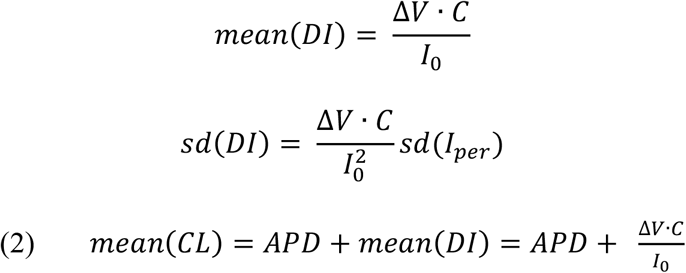

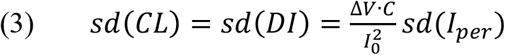

To derive a fundamental relationship between *mean(CL)* and *sd(CL)*, we first express *1/I*_*0*_ from (3) in terms of *sd(CL)*

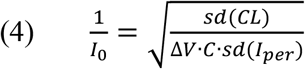

and then we substitute for *1/I*_*0*_ using (4) in equation (2) obtaining a square root relationship:

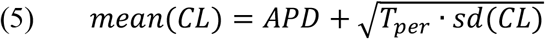

where *T*_*per*_ is perturbation time given as

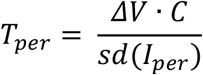

In terms of *mean(HR)* equation (5) yields the inverse squire relationships stated in the paper tittle:

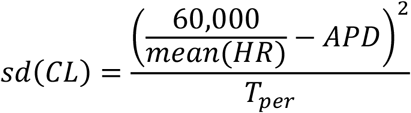

for *HR* in bpm and *APD* and *T*_*per*_ in ms (see respective data fit in Fig.1C), or in basic terms:

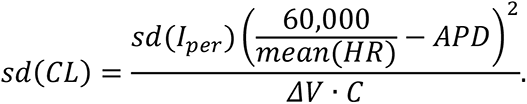

## Acknowledgments

This work was supported by the Intramural Research Program of the NIH, National Institute on Aging. A.V.M. acknowledges the support of the Royal Society University Research Fellowship UF160569. A.V.M. also thanks Prof. Boyett at the University of Manchester for his hospitality and related discussions.

